# Genomic Diversity Illuminates the Environmental Adaptation of *Drosophila suzukii*

**DOI:** 10.1101/2023.07.03.547576

**Authors:** Siyuan Feng, Samuel P. DeGrey, Christelle Guédot, Sean D. Schoville, John E. Pool

## Abstract

Biological invasions carry substantial practical and scientific importance, and represent natural evolutionary experiments on contemporary timescales. Here, we investigated genomic diversity and environmental adaptation of the crop pest *Drosophila suzukii* using whole-genome sequencing data and environmental metadata for 29 population samples from its native and invasive range. Through a multifaceted analysis of this population genomic data, we increase our understanding of the *D. suzukii* genome, its diversity and its evolution, and we identify an appropriate genotype-environment association pipeline for our data set. Using this approach, we detect genetic signals of local adaptation associated with nine distinct environmental factors related to altitude, wind speed, precipitation, temperature, and human land use. We uncover unique functional signatures for each environmental variable, such as a prevalence of cuticular genes associated with annual precipitation. We also infer biological commonalities in the adaptation to diverse selective pressures, particularly in terms of the apparent contribution of nervous system evolution to enriched processes (ranging from neuron development to circadian behavior) and to top genes associated with all nine environmental variables. Our findings therefore depict a finer-scale adaptive landscape underlying the rapid invasion success of this agronomically important species.

## Introduction

One of the main goals of ecological and evolutionary genomics is to understand how organisms evolve in response to novel environments. Biological invasions, while often ecologically and economically damaging, represent unique opportunities to build our understanding of local adaptation, as natural experiments that expose introduced species to new biotic and abiotic factors on contemporary time scales (Lee 2002; Prentis et al. 2008; Colautti and Lau 2016). Invasive species can exhibit rapid phenotypic and genetic changes during the invasion process, driven by various evolutionary mechanisms such as selection, drift, mutation, and gene flow (Colautti and Lau 2016; Hodgins et al. 2018). These changes can result in the adaptive evolution of invasive populations to the novel environments they encounter (Colautti and Barrett 2013). Although there have been emerging studies on the evolutionary biology of invasive species in recent years, the source and nature of the genetic variation underlying such adaptation are still not well characterized (Reznick et al. 2019; Welles and Dlugosch 2019).

*Drosophila suzukii*, also known as spotted wing *Drosophila*, is a promising model for studying adaptive evolution during invasions. *Drosophila suzukii* is a highly polyphagous vinegar fly that originated from Asia (Peng 1937; Kanzawa 1939; Tan et al. 1949). It first expanded to Hawaii in 1980, and within the past 15 years, it has invaded North America and Europe, and subsequently Réunion Island (Indian Ocean) and South America (Asplen et al. 2015), then northern and sub-Saharan Africa (Boughdad et al. 2021; Kwadha et al. 2021). *Drosophila suzukii* differs from other *Drosophila* species in its unique ability to oviposit on both unripe and ripe fruits, using its serrated ovipositor to pierce the skin of soft-skinned fruits. This has allowed it to exploit a novel ecological niche and avoid competition with other vinegar flies that typically feed on overripe and rotting fruits (Cini et al. 2012; Atallah et al. 2014), causing severe economic losses to fruit crops (Knapp et al. 2021). It also exhibits remarkable genetic diversity and phenotypic plasticity in behavior, morphology, and physiology, e.g., temperature and desiccation tolerance (Little et al. 2020; Olazcuaga et al. 2020), which may notably facilitate its adaptation to different climatic conditions and host plants (Gibert et al. 2019; Little et al. 2020). To obtain a comprehensive evolutionary genetic understanding of the invasion success of *D. suzukii*, we need to understand the genetic basis and ecological drivers of adaptive evolutionary changes that have allowed this species to occupy diverse worldwide environments.

Multiple genetic studies have investigated the demographic history of invasive populations of *D. suzukii*. Such studies have found greater levels of genetic structure between than within continents (Adrion et al. 2014; Lewald et al. 2021), suggesting independent invasions into Europe and North America. These inferences were supported by an approximate Bayesian analysis of microsatellite data, which also indicated that some invading populations had multiple genetic sources (Fraimout et al. 2017). While minor differences are suggested in the specific admixture events that have occurred (Fraimout et al. 2017; Lewald et al. 2021), and inferences of the geographic origins of invading populations are limited by incomplete sampling from the species’ Asian range, these studies suggest some emerging consensus about the invasion history of *D. suzukii*.

In contrast, the genetic basis of environmental adaptation in *D. suzukii* is still largely unexplored. Among the few relevant studies is that of Olazcuaga et al. (2020), which used genomic sequencing data from 22 worldwide populations to search for SNPs with greater frequency differences between Asian (China, Japan) and non-Asian populations than are observed at most loci due to founder event bottlenecks, in hopes of identifying genetic variants that may underlie the invasion success of introduced populations. A subsequent study examining this same data set also found a small number of transposon insertions with strong frequency differences between American/European and Asian populations (Mérel et al. 2021). Another study identified *FST* outliers among Hawaiian *D. suzukii* populations (Koch et al. 2020). Apart from these limited comparisons, the genetic changes that may have helped *D. suzukii* to adapt to specific environmental changes are entirely unknown.

With the increasing availability of multi-population genomic resources, genotype-environment association (GEA), also known as environmental association analysis (EAA), is becoming a widely-used approach to understand the potential relationship between specific environmental factors and adaptive genetic variation (Rellstab et al. 2015). GEA is also useful in identifying subtle changes in allele frequencies that are difficult to detect with outlier tests based on traditional population genomic approaches, especially when the number of studied populations is relatively large, and there is high gene flow counteracting patterns of local adaptation (Kawecki and Ebert 2004). The capability of GEA to identify adaptive genetic changes and environmental drivers of local adaptation has been demonstrated with whole-genome pool-seq data from *Drosophila melanogaster* (Bogaerts-Márquez et al. 2021). Therefore, GEA could be helpful in understanding the genotype-environment relationships underlying the invasion success of *D. suzukii*. Performing GEA on this model system could also facilitate insights into environment-driven genetic changes during rapid biological invasions, as well as mechanisms driving insect adaptation to rapid climate shifts *in situ*.

In the present study, we perform population genomic analyses on whole-genome pool-seq data of 29 population samples from native and invasive ranges to investigate the environmental adaptation of *D. suzukii*. We investigate the geographic pattern of genetic diversity and population structure, in part to inform our choice of GEA methodology. We then test the association between SNP frequencies and nine environmental variables across sampling locations, identifying both specific and shared functional signatures of adaptation to these diverse selective pressures.

## Results

### Genomic Diversity and Population Structure of *D. suzukii*

To investigate the genetic diversity of *D. suzukii* and to examine the genetic input for environmental association analysis, we summarized the genomic polymorphism of 29 *D. suzukii* populations derived from Asia (n=8), Europe (n = 11), and North and South America (n = 10), encompassing both newly reported and previously published samples (Figure 1A; Table S1; Olazcuaga et al. 2020). Whole-genome sequences were obtained from 29 pooled samples consisting of 50-212 female and male individuals. The depth of mapped reads after quality control ranged from 23X to 66X among population samples, with an average of 45X (Table S1).

**Figure 1.**
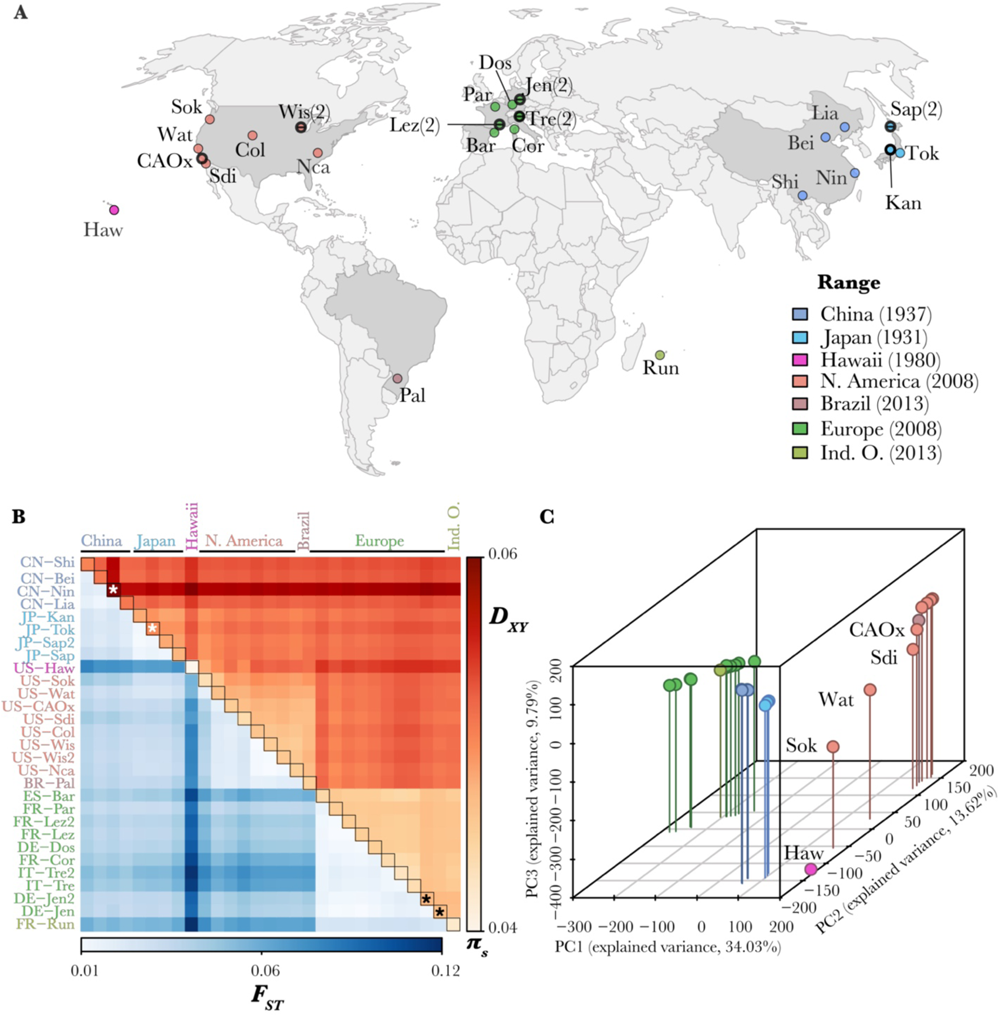
*Drosophila suzukii* populations show maximal diversity in eastern Asia and continent-level genetic structure. (A) The geographic locations of the studied 29 natural populations are depicted as dots. In addition to the 22 populations sampled by Olazcuaga et al. (2020), populations newly sampled at independent locations are circled in black. Populations newly sampled at nearby locations are circled and center-dashed in black, with the number of total population samples in brackets. The year of the first recorded occurrence in each geographic range (colored grey in the map) is given in brackets in the color legend. Further information about each sample is presented in Table S1. (B) Population differentiation in allele frequencies (*F_ST_*; lower triangle), between-population sequence distances (*D_XY_*; higher triangle), and within-population nucleotide diversity (*π_S_*; diagonal) across autosomal synonymous SNPs are displayed as a heatmap. Population names are colored by their geographic region. Asterisks indicate samples contaminated by other *Drosophila* species, which may affect estimation of these statistics. (C) Autosomal genetic structure is shown by three-dimensional principal components analysis (PCA) based on allele frequencies of the two most frequent alleles across all populations. Each dot represents a population. Labeled are Hawaii and western coastal US populations. See the X chromosome version of (B) and (C) in Figure S2.

We first estimated nucleotide diversity (*π_S_*) across synonymous SNPs to investigate the effects of rapid invasions on neutral genetic diversity. The genome-wide *π_S_* ranged from averages of 0.045 (autosomes) and 0.025 (X chromosome) in introduced populations, to averages of 0.051 (autosomes) and 0.034 (X chromosome) in Asian populations (Table S2). Previous studies have hypothesized a relatively wide native range of *D. suzukii* in East and Southeast Asia (Adrion et al. 2014; Fraimout et al. 2017), and the acute drop in *π_S_* of introduced European and American populations relative to that of the native Chinese and Japanese populations reflects previously reported founder event bottlenecks (Figure 1B, diagonal; Figure S1, S2A). The observed patterns were also recapitulated with SNPs at all types of sites (Figure S3). Lower *π_S_*and more contrasting among-population *π_S_* differences in X chromosome reflect effectively prolonged bottlenecks due to their lower effective population size (Pool & Nielsen, 2007; Figure S1). We also observed a greater loss of rare alleles in the invasive populations as a typical consequence of bottlenecks (Figure S4). These founder events also increased genetic differentiation as measured by *F_ST_*, especially between continents (Figure 1B; Figure S3), and particularly for the X chromosome (Figure S2A).

Subsequent to our analysis, contamination in three of the published samples was reported, involving reads from the closely-related *D. subpulchrella* and from *D. immigrans* (Gautier 2023), which can be estimated but not fully removed. The estimated level of *D. subpulchrella*-sourced contamination for Tokyo, Japan (JP-Tok) was reduced from 4.47% to 3.58% after our alignment and quality filtering pipeline (Table S3), which is slightly higher than the baseline noise level of 1.14% to 2.78% from samples known to be pure *D. suzukii* (Gautier 2023). The published DE-Jen sample’s contamination from *D. immigrans* was reduced from 5.79% to 0.35%. Our unpublished sample from the same area showed 0.28% *D. immigrans* contamination from mapped reads, whereas the other unpublished samples showed no evidence for contamination (Table S3). However, a higher proportion of reads from *D. subpulchrella* remained in the east Asian sample CN-Nin (10.38%, versus 14.95% from raw reads). Hence, it is likely that the elevated *π_S_* of CN-Nin (and likewise its elevated *D_XY_*values from the analysis described below) reflect an artefact of this contamination.

We next analyzed genetic structure among the sampled populations, using the summary statistics *D_XY_* and *F_ST_*, and via principal component analysis (PCA) of population allele frequencies, particularly since the pattern of genetic structure present may influence the performance of GEA methods (Rellstab et al. 2015). The top three principal components (PC 1-3) explained 57.44% (autosomes, Figure 1C) and 72.35% (X chromosome, Figure S2B) of the variance among populations. In both autosomes and X, three-dimensional PCA and matrices of both *D_XY_* and window *F_ST_* recapitulated both continuous and hierarchical geographic structure, which must be accounted for in GEA. These results together showed the expected clustering of populations into four distinct ranges (East Asia, Hawaii, Americas, and Europe; Figure 1B, 1C; Figure S2). Much of the observed population differentiation is most likely due to founder event bottlenecks and admixture during worldwide expansion (Fraimout et al. 2017), whereas migration following population establishment would need to be overwhelmingly high to have significant impacts given the very brief time scale of the global invasion. The two more northerly populations from the western US, Oregon (US-Sok) and central California (US-Wat), show an affinity with the Hawaiian population, which aligns with the suggestion that these populations received a genetic contribution from Hawaii in addition to East Asia (Fraimout et al. 2017), while populations from southern California and the central and eastern US show less evidence of such admixture.

Our focus above on diversity and differentiation at synonymous sites was motivated by the low selective constraint expected at these sites, as confirmed by our analysis of divergence between *D. suzukii* and its relative *D. biarmipes* (Ometto et al. 2013; Suvorov et al. 2022; Figure S5). We also note that compared to the divergence between *D. melanogaster* and *D. simulans* (Begun et al. 2007; Lange and Pool 2018), *D. suzukii* and *D. biarmipes* show relatively higher ratios intergenic (or intronic) divergence to non-synonymous divergence (Figure S5). Compared with *D. melanogaster*, the genome of *D. suzukii* contains a notable expansion of repetitive sequences (Paris et al. 2020), which could reflect a lower long-term effective population size (*N_e_*) in the *D. suzukii* lineage. However, the greater *π* of *D. suzukii* than *D. melanogaster* (Figure S1; Lack et al. 2016a; Lewald et al. 2021) could instead indicate a greater *N_e_* for *D. suzukii* within the past 4*N_e_* generations.

### Polymorphism and the Genomic Locations of *D. suzukii* Contigs

If extensive genomic regions of low recombination are present in a genome-wide scan for local adaptation, then because of the larger-scale influence of natural selection on linked sites in such regions (Smith and Haigh 1974; Charlesworth et al. 1993), the precision of outlier identification will be reduced (Lotterhos and Whitlock 2015; François et al. 2016). Although we do not have recombination rate estimates for *D. suzukii*, we can begin to assess the genomic abundance of low recombination regions through an examination of nucleotide diversity, in light of its expected correlation with recombination rate (Begun and Aquadro 1992). Fortunately, we observe that regions of low nucleotide diversity (which probably coincide with regions of low recombination) cover a relatively small fraction of the genome (Figure 2). These patterns are more similar to those in *D. simulans* than to *D. melanogaster* – which has broader centromeric regions of low crossing over on the autosomes (Langley et al. 2012) – potentially suggesting a relatively weaker suppression of crossing over in the centromere-proximal regions in *D. suzukii*.

**Figure 2.**
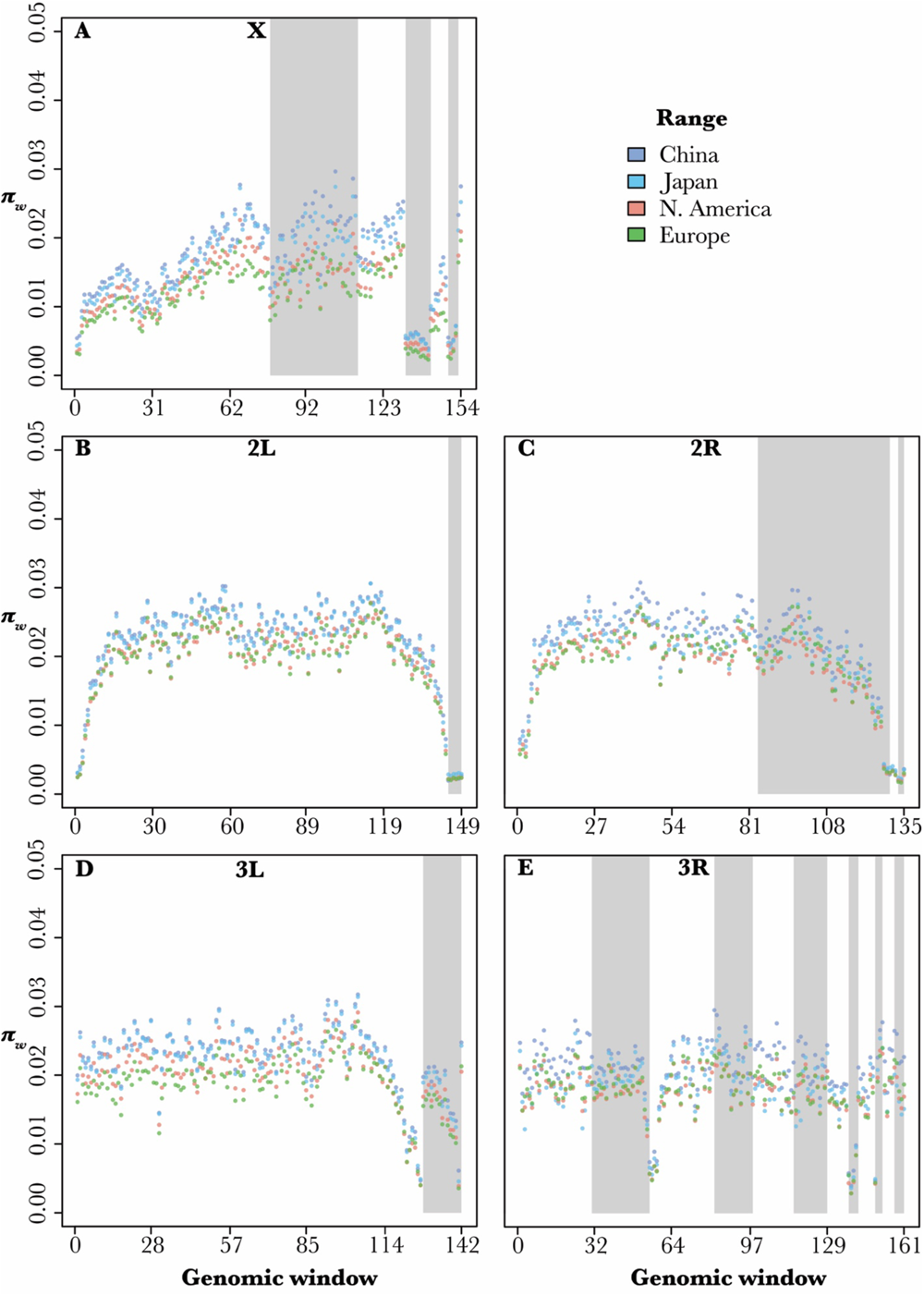
Chromosomal distribution of genetic polymorphism in *D. suzukii* informs the ordering and orientations of contigs, as well as levels of centromeric and telomeric repression. Window nucleotide diversity (*π*_+_) values are displayed across (A) the X chromosome and major autosomal arms (B) 2L, (C) 2R, (D) 3L, (E) 3R. Chromosome 4 is not shown as it only contains 12 windows. Each window is a continuous genomic region that includes 125,000 analyzed sites. Each dot represents the average *π*_+_across populations within their geographic range as colored. Only populations from major continental ranges are shown for chromosomal patterns to be clear. Within each chromosome arm, contigs are ordered by length. Separate contigs are indicated by grey or white shading.

Population genomic data may also provide clues regarding genome structure. The *D. suzukii* reference assembly (Paris et al. 2020) is somewhat fragmented, with two or more contigs assigned to each major chromosome arm, but with unknown orientation with respect to each other. In other examined *D. melanogaster* group species, crossing-over rate and window nucleotide diversity (*π*_+_) are reduced in centromere-and telomere-proximal regions (*e.g.,* True et al. 1996). When we examined the chromosomal distribution of *π*_+_ (Figure 2), we noticed that certain arrangements of contigs would result in the expected patterns of reduced diversity at the ends of each arm, and relatively smooth shifts in the diversity of large windows. For instance, the second longest contig in chromosome arm 3L is likely to the left of the longest contig in a reverse orientation, potentially with the smaller high diversity contig in between, (Figure 2D), although either of the low diversity contig end could be centromeric or telomeric. Similarly in chromosome arm 3R, although the assembly was more fragmented and the centromere-or telomere-proximal regions consist of multiple contigs, the second and third longest contigs appeared to contain the transitions between the high diversity mid-arm region and the centromeric/telomeric zones (Figure 2E). Therefore, genome-wide polymorphisms could serve as useful information to aid the ordering and orienting of contig-level genome assemblies (Nakabayashi and Morishita 2020), although a combination of both systematic neighbor-matching approaches and manual correction would be needed, and integration with other forms of evidence (*e.g.,* comparative genomic or experimental data) would be preferable.

We next leveraged our large pooled sequencing data set to improve inferences about which contigs map to the X chromosome, in part so that X-linked and autosomal contigs could be more accurately partitioned in our subsequent GEA. Out of a total of 546 contigs, 313 were previously assigned to autosomes and X chromosome through either direct mapping or comparing a female-to-male read depth ratio (Paris et al. 2020). We added to these annotations by implementing an approach based on correlations in sequencing depth of coverage across population samples that included varying numbers of females and males (see Materials and Methods). Based on this analysis, we assigned 170 contigs as autosomal or X-linked (Table S4). Our classification of previously-assigned contigs was 96% consistent with past inferences, but we corrected four previous assignments that were based on female-to-male read depth ratio, whereas our method did not assign eight previously-assigned contigs.

### Selection of Robust Methodology and Distinct Environmental Variables for GEA

The worldwide expansion of *D. suzukii* has exposed this species to selection pressures from varying local environmental conditions (Olazcuaga et al. 2020; Mérel et al. 2021). To identify environmental factors that have contributed to adaptive genetic differentiation at various levels and loci under positive selection, we performed a whole-genome scan using GEA analysis between environmental and genetic differentiation (see Materials and Methods). We focused on the method BayeScEnv because it has been reported to produce fewer false positives in the presence of hierarchical genetic structure such as the continental-scale patterns documented here (de Villemereuil and Gaggiotti 2015). We excluded SNPs with a global average minor allele frequency (MAF) below 5%, which should minimize the influence of the contamination documented above. No alleles specific to contaminating species should meet that threshold, so the only remaining effect should be a slight bias in allele frequency estimation toward ancestral variants in contaminated samples at genuine *D. suzukii* SNPs. Given the low levels of contamination present and the small number of samples affected, this effect may represent a modest source of noise, but it should not lead to spurious adaptation signals.

Our selection of environmental variables for GEA started with a preliminary set of 26 candidate variables that are potentially relevant in the adaptation process of *D. suzukii* (Figure 3A), based on data availability and prior knowledge of *Drosophila* ecology (Kellermann et al. 2012; Hamby et al. 2016; Bogaerts-Márquez et al. 2021). Among common strategies of performing GEA with a large set of relevant environmental variables, univariate association with all environmental variables could increase the number of statistical tests, thus increasing the difficulty of controlling rates of false discovery. On the other hand, including multiple variables might cause a multicollinearity issue (Rellstab et al. 2015). We opted to retain nine of the least correlated environmental variables for univariate tests, representing altitude, wind speed, as well as multiple aspects of temperature, precipitation, and human land usage for GEA analysis (Figure 3B). Although the temperature of the coldest quarter had a significant negative correlation with wind speed, we kept both variables for GEA, as cold stress and wind-related factors are known to be potential drivers of local adaptation in *Drosophila* (Bogaerts-Márquez et al. 2021). As indicated by their coefficients of variation (*CVs*), the selected environmental variables had moderate (15% ≤ *CV* < 30%) to high (*CV* ≥ 30%) variability across our sampling locations (Table S4).

**Figure 3.**
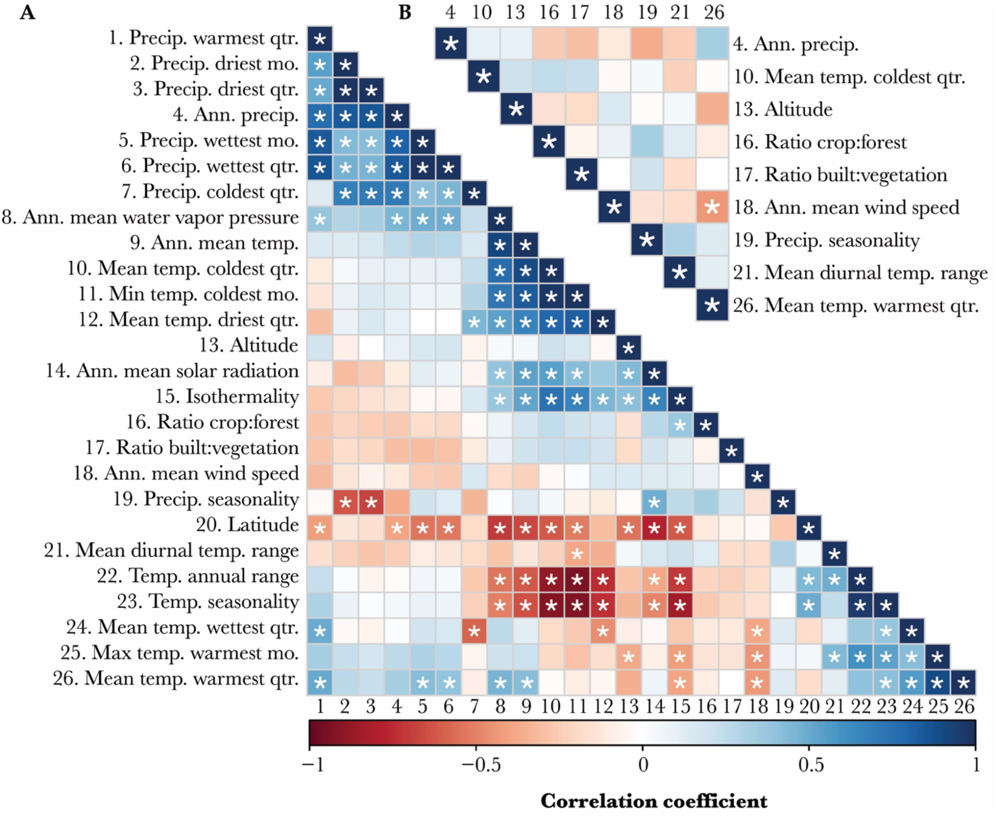
Identification of least-correlated environmental variables for genotype-environment association (GEA) analysis in *D. suzukii*. (A) Pairwise correlations among a preliminary set of 26 environmental variables that are potentially impactful on *D. suzukii*. (B) A final set of nine of the most relevant and least correlated environmental variables that were chosen for GEA analysis. The Pearson correlation coefficients are colored from -1 (perfect negative correlation) to 1 (perfect positive correlation). Significance correlations (p < 0.05) are indicated by asterisks. See Table S4 for environmental values used to calculate correlation coefficients.

### Widespread Signals of Recent Adaptation to Diverse Environments

From 5,752,156 genome-wide SNPs with a global average MAF higher than 5%, we identified an average of 3,033 (SD = 823.4) unique candidate variants that were significantly (genome-wide *q* < 0.05) associated with each of the nine candidate environmental variables (Table S6). These variants corresponded to an average of 3,345 overlapping or neighboring genes (Table S6), suggesting that selection pressures from the tested environmental variables (or correlated factors) have meaningfully contributed to adaptive genetic variation in *D. suzukii*, even on the brief time scale of its worldwide expansion. Among all tested environmental factors, mean temperature of the coldest quarter was associated with the greatest number of putatively adaptive variants (4,250 SNPs). Two precipitation-related variables, annual precipitation (4,141 SNPs) and precipitation seasonality (3,389 SNPs), have the next largest loci count. The ratio of built area to vegetation (i.e., crops and forests) and the ratio of crops to forests were associated with the fewest genetic variants (2,369 and 1,608 SNPs, respectively).

To reveal the potential genetic and functional basis of invasion success under multiple environmental challenges throughout the species’ range, we first examined the functions of genes linked to the top 10 environment-associated loci for each variable as ranked by association q-value, and then by the *g* parameter estimating the sensitivity to environmental differentiation as a tie-breaker. We found many of these genes have known functions that could facilitate adaptation to the associated environmental factor (Table S7).

Among genes linked to altitude-associated loci, the second-ranked candidate *ab* is known to control wing size in *Drosophila* (Simoes da Silva et al. 2019). Interestingly, wing size was found to have increased in a highland Ethiopia *D. melanogaster* population, potentially assisting flight in thin, cool air (Lack et al. 2016b). Ranked next to *ab* is *Gbs-70E*, which plays roles in glycogen metabolism and the development of eggs inside the maternal ovary (Kerekes et al. 2014).

Another top gene, the lysine demethylase *Kdm2*, is upregulated in response to hypoxia (Batie et al. 2017).

With wind speed, the top first candidate *Ttc30* is an essential gene in the biogenesis of sensory cilia, which are key to both chemosensory and mechanosensory functions in *Drosophila* (Avidor-Reiss et al. 2004; Avidor-Reiss and Leroux 2015). Another top candidate, *Arr2*, is involved in olfaction, hearing, and vision (Alloway and Dolph 1999; Elaine Merrill et al. 2005; Senthilan et al. 2012). In light of the relevance of wind for insect flight, we also noted that a third top candidate, *vn*, is a developmental gene named for its wing phenotype (Wang et al. 2000).

There is also some evidence for precipitation-related local adaptation. The top gene *mmy* associated with precipitation seasonality (i.e., annual range of precipitation) was shown to regulate chitin synthesis and cuticle production. Since precipitation is correlated with desiccation resistance across the *Drosophila* phylogeny (Kellermann et al. 2012), *D. suzukii* may have developed adaptive strategies of modifying chitin biosynthesis under conditions of desiccation (Rezende et al. 2008; Clark et al. 2009), which was also implied in seasonal plasticity of natural *Drosophila* populations (Shearer et al. 2016; Horváth et al. 2023). In addition, the gene *osy* (*CG33970*) contributes to the formation of the outer cuticle layer and is expressed more highly in *D. suzukii* than in *D. melanogaster* (Wang et al. 2020). Furthermore, two of the top genes associated with annual precipitation (*Abd-B* and *bab1*) regulate cuticle pigmentation (Rogers et al. 2013), which may or may not correlate with desiccation tolerance in *Drosophila* species (Wang et al. 2021). We also note that although environmental fitness effects on these testes-expressed genes are not known, the same SNP near *CG17944* and *nxf4* was among the highest-scoring variants for both annual precipitation and precipitation seasonality (variables that have a non-significantly negative correlation between them; Figure 3).

Another important environmental barrier to invasion success is temperature. For the mean temperature of the coldest quarter, a top gene was *Ac78C*, which has roles in circadian regulation and taste (Ueno and Kidokoro 2008; Duvall and Taghert 2013). With the mean temperature of the warmest quarter, the top genes *crp*, *Mrtf*, and *Ubx* help control the development of trachea (Han et al. 2004; Guha and Kornberg 2005; Wong et al. 2015), which may be important in limiting water loss in hot environments (Gibbs et al. 2003).

For the ratio of built to vegetated area, a different variant near the cuticle-related gene *osy* (which was also indicated above for precipitation seasonality) was detected. Another top outlier was the nervous system gene *trv*, which is involved in thermosensitivity (Honjo et al. 2016). For the relative levels of crop and forest cover, the first-ranked variant was near *Mtk*, which encodes an antifungal and antibacterial peptide (Levashina et al. 1995), and we note that mushrooms (which are more available in forest) have been proposed as overwintering food sources for *D. suzukii* (Wallingford et al. 2018), and the evolution of immune genes has been found to differ strongly between mushroom-feeding and human commensal *Drosophila* species (Hill et al. 2019). With regard to the differential light environments entailed by forest versus farm habitats, we note that the next highest gene, *CadN2*, helps connect photoreceptor neurons to their targets (Prakash et al. 2005).

Beyond the top candidate genes that have related functions to specific types of environmental changes, we also found a wide range of nervous system genes associated with multiple environmental factors. For instance, among the top five altitude-associated loci, three have known functions in the nervous system of *Drosophila*, including the first-ranked gene *Cmpy*, which enables proper growth control at neuromuscular junctions (James and Broihier 2011), *ab*, which regulates dendritic complexity (Li et al. 2004; Sugimura et al. 2004), and *not*, which is essential for stabilizing synaptic homeostasis within glia (Wang et al. 2020). Such genes were also linked to top-10 loci associated with wind speed (*dpr6*), precipitation (*Msp300*, *Tusp*, *5-HT2A*), temperature (*ATP6AP2*, *CG13579*, *D*), and land use variables (*Bsg*, *Mp*, *velo*). Different variants associated with *scrt*, a regulator of neuronal cell fate, were among the top results for both mean diurnal range and the ratio of crop to forest cover.

### Functional Commonalities in the Adaptation to Diverse Selective Pressures

Next, we examined environment-specific adaptation on a more comprehensive basis through a gene ontology (GO) enrichment analysis of the top 500 genes associated with each environmental variable (Figure 4). As correlates of temperature in the coldest quarter, cAMP metabolic process was the top enriched category, followed by two other related purine metabolism groupings. We note that cAMP is important in circadian regulation (*e.g.*, Palacios-Muñoz & Ewer, 2018), which is known to play an important role in *Drosophila* environmental adaptation (*e.g.,* Helfrich-Förster et al. 2020), as also implicated by the presence of “entrainment of circadian clock” on our top GO category list for altitude. More broadly, purine metabolism was inferred as a strategy of cold acclimation in *D. suzukii* (Enriquez and Colinet 2019). For diurnal temperature range, the top category was “regulation of growth”, and we note that some drosophilids have evolved to have larger body sizes in more challenging thermal environments (Gilchrist and Partridge 1999; Calboli et al. 2003; Lack, Yassin, et al. 2016).

**Figure 4.**
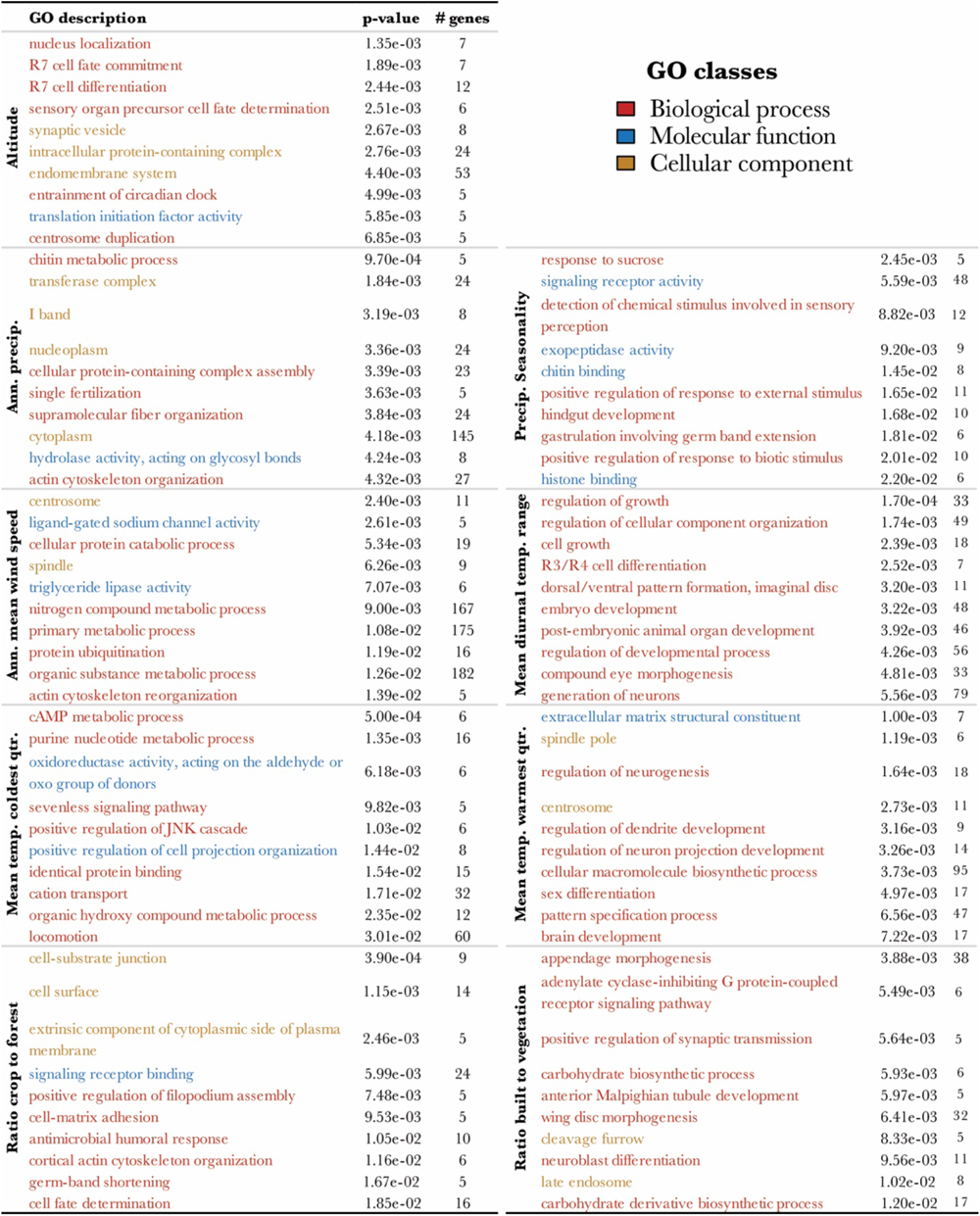
GO enrichment analysis of candidate genes from the gene-environment association analysis of *D. suzukii*. The top 10 GO categories enriched by the top 500 genes associated with each environmental variable are shown in each panel (labelled on the left), with permutation p-values and the number of associated genes in each GO category. Descriptions of GO categories are colored by their GO class (see legend at top right). Only GO categories including more than five associated genes are listed here. For a full list of enriched GO categories, see Table S8.

With precipitation, we identified “chitin metabolic process” as the top GO term associated with annual precipitation, as well as “chitin-binding” with precipitation seasonality. Together with the chitin synthesis genes we described above for precipitation, adaptation to the overall intensity and seasonal variation of precipitation by modifying cuticular chitin may be implied. For crop to forest ratio, the category “antimicrobial humoral response” included the top gene *Mtk* listed above.

As broader evidence for a shared (or biologically similar) underlying genetic basis of adaptation to multiple environmental factors, we examined the overlap of the most significant genes and most enriched GO categories between different environmental variables. We found the top candidate genes to be mostly associated with both temperature and precipitation. The gene sets showed relatively greater overlap among climatic variables (including altitude), whereas the two land usage variables had less overlap with climatic variables or with each other (Figure 5A). Since the patterns of shared genes cannot be fully explained by correlations between environmental values (Figure 3B), at least some of the genes may have been responding to multiple selective pressures. The overall proportions of shared GO categories were lower than those of shared genes, indicating that the shared genes do not necessarily lead shared functional categories between environmental variables. Relatively higher GO term sharing was observed between altitude and either wind speed or precipitation, and between diurnal temperature range and temperature of the warmest quarter (Figure 5B). Based on the shared genes and GO terms observed, it is possible that during the rapid range expansion of *D. suzukii*, pleiotropy may have facilitated local adaptation to multiple selective pressures (Hämälä et al. 2020; Kinsler et al. 2020).

**Figure 5.**
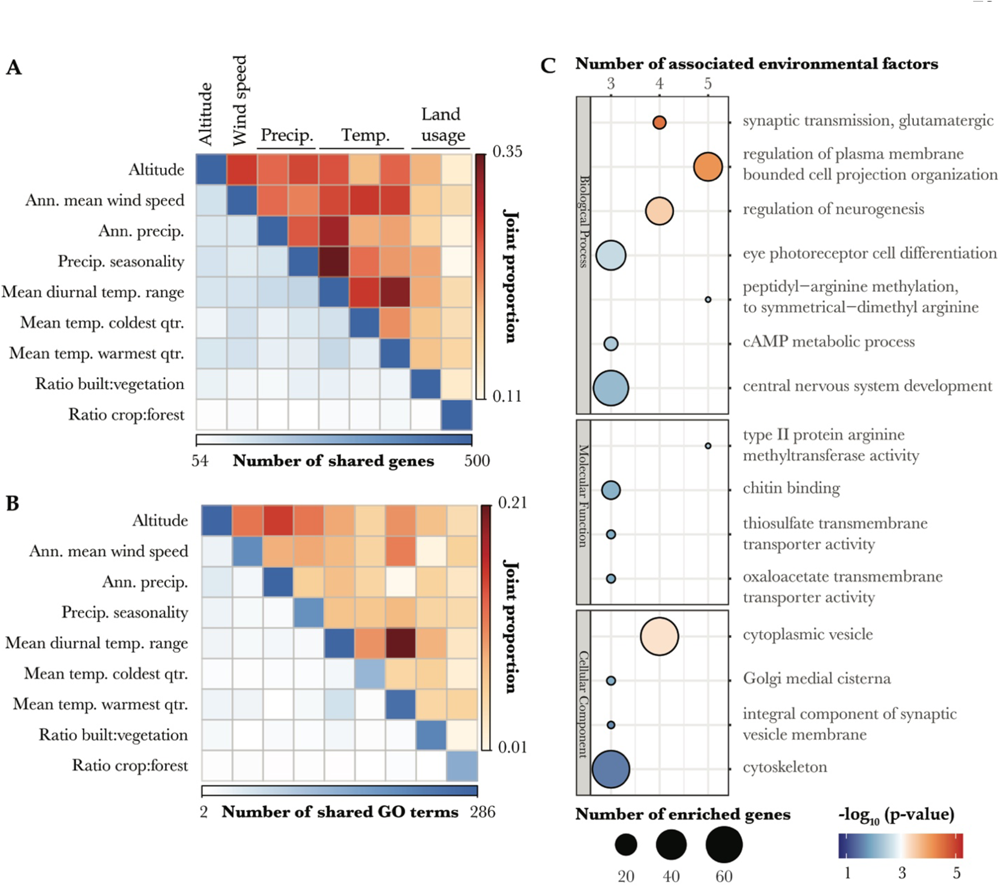
Overlapping genes and GO categories among environmental factors reveal the shared genetic and functional basis of environmental adaptation in *D. suzukii*. The numbers and proportions of shared (A) environment-associated genes and (B) enriched GO categories among environmental factors are shown in heatmaps. Here, joint proportion represents the fraction of the genes or GO terms associated with either of two environmental variables that are associated with both variables. (C) Top GO categories of each type are depicted as bubbles. Bubbles are colored by the negative logarithm of the combined p-value of enrichment across all environmental variables, and are scaled by the number of enriched genes. The number of environmental variables that enrich a given GO category is indicated by the top horizontal axis.

Consistent with our gene-based analyses of universal adaptive function, we found three of the top shared biological processes clearly related to nervous system functions, including the topmost “synaptic transmission, glutamatergic” (shared by altitude, wind speed, diurnal temperature range, and temperature of the warmest quarter), “regulation of neurogenesis” (altitude, diurnal temperature range, temperature of the warmest quarter, and ratio of built area to vegetation), and “central nervous system development” (precipitation seasonality, diurnal temperature range, and temperature of the warmest quarter) (Figure 5C). Further, as mentioned above, “cAMP metabolic process” (shared by temperature of the coldest quarter, precipitation seasonality and ratio of built area to vegetation) could entail neurologically modulated changes in circadian behavior. Each of these traits was associated with at least three environmental variables, which suggests a multifaceted role for nervous system evolution in facilitating the invasion success of *D. suzukii* under multiple environmental challenges.

## Discussion

We performed population genomic analyses of 29 population samples of *D. suzukii* to investigate the genomic diversity and environmental adaptation of this highly invasive species across its worldwide distribution. Our data supported a genetic grouping of these populations into four primary geographic regions: eastern Asia (containing the native range), Hawaii, the Americas, and Europe. We also confirmed that all non-Asian populations have reduced diversity, consistent with moderate founder event bottlenecks in introduced populations.

Our analyses also added to our knowledge of the *D. suzukii* genome and its evolution. We used population genomic data to improve the classification of X-linked and autosomal contigs. We determined that relatively few contigs showed strongly reduced nucleotide diversity, implying that only a small fraction of the genome experiences minimal crossing-over. And we documented the influence of an expanded repeatome on non-coding divergence.

The above analyses placed us in a more confident position to perform a robust analysis of genotype-environment association (GEA). We selected nine distinct environmental variables, including altitude, wind speed, precipitation, temperature, and human land usage. Our results suggested extensive local adaptation in response to specific environmental challenges, along with appreciable sharing of genes and functional pathways underlying invasion success across multiple environmental pressures, which were most obvious with nervous system genes.

### Environmental Drivers of Adaptation in *D. suzukii*

Here, we presented a GEA analysis that investigated the most geographically and genetically diverse set of *D. suzukii* populations and the most comprehensive set of environmental factors to date, which enabled unprecedented power to capture even minor adaptive genetic differentiation in response to distinct environmental challenges during the species’ rapid invasions. In addition to our identification of climatic factors including temperature, precipitation-related variables and wind speed as the most frequently correlated with putatively local adaptative variants (consistent with previous GEA analysis in *D. melanogaster*, *e.g.,* Bogaerts-Márquez et al. 2021), we also for the first time identified large numbers of genome-wide variants associated with altitude and human land usage-related variables (Table S6), which were not included in most GEA studies despite their potential significance to local adaptation. In particular, the detected associations with ratios of developed land to vegetation and of cropland to forests highlights the ecological impacts of urbanization and agriculture on natural populations of insects. Since the prior selection of environmental variables is critical for successful GEA analyses, we also provided an instructive example for correlation-based selection to identify the most relevant and least redundant environmental factors (Rellstab et al. 2015).

### Nervous System Evolution is Ubiquitous in Environmental Adaptation of ***Drosophila***

In *D. suzukii*, we found nervous system and related sensory and behavior annotations associated with top genes for all nine environmental variables studied. Concordantly, we found that GO categories related to the nervous system were among the most shared across environmental variables (Figure 5C). In *D. melanogaster*, related GO categories like ‘neuron development’, ‘nervous system development’ and ‘eye development’ were also enriched among genes associated with environmental variation among natural populations within North America or Europe, and across seasons within Europe (Bogaerts-Márquez et al. 2021). GO categories associated with the nervous system have also shown evidence of positive selection in various genome scans of *D. melanogaster* (Langley et al. 2012; Pool et al. 2012; Pool 2015), including a study of parallel evolution in cold-adapted populations (Pool et al. 2017). Given the morphological evidence of neuron-muscular junction evolution across the entire *Drosophila* phylogeny (Campbell and Ganetzky 2012), we therefore propose a broad adaptive importance of the nervous system in *Drosophila* species and potentially other insects. Such evolutionary processes may have either maintained ancestral neural functions in novel challenging environments, or created novel phenotypes that better fit the new optima arising from complex combinations of environmental factors.

### Cautions with Interpretations of Association Results and Future Directions

While we have generated intriguing hypotheses about gene functions that may underlie the environmental adaptation of *D. suzukii*, it is difficult to distinguish between correlated environmental selective pressures that may have driven the detected associations, including not only the 17 environmental factors that were excluded in the process of variable reduction, but also correlated biotic or abiotic factors not represented in global databases. As an intrinsic limitation of GEA analysis that cannot be accounted for by applying stricter thresholds, associations observed with a particular environmental factor might stem from adaptation to other covarying factors (Rellstab et al. 2015). For example, the two tracheal branching genes *crp* and *Mrtf* (Han et al. 2004; Wong et al. 2015) associated with mean temperature of the warmest quarter could represent adaptations to reduce water loss under conditions of elevated water vapor pressure (Telonis-Scott et al. 2012), which is closely related to humidity and has a significant positive correlation with mean temperature of the warmest quarter (Figure 3B).

Therefore, expanded characterization of the relationships between genotype, phenotype, and fitness in this species is needed to further clarify the functional and phenotypic interpretations associated with certain environmental factors and genes. Experimental validations that leverage RNA interference (Boutros and Ahringer 2008) and/or transgenic overexpression (Prelich 2012) to modify the expression of associated genes, and/or genome editing techniques (Stern 2014; Turner 2014; Shalem et al. 2015) to target putatively adaptive variants would also bring a more solid understanding about the invasive biology of this species in distinct environments. Such functional studies could be complemented by population experiments under controlled laboratory environments or field conditions (*e.g.,* Behrman et al. 2015; Rudman et al. 2022), in order to more clearly demonstrate the connections between specific selective pressures and alleles or traits of interest.

### Broader Impacts and Significance

Our work integrates genetic and environmental data to improve the reconstruction of the invasion genomics of a crop pest carrying significant economic costs (Knapp et al. 2021), which will hopefully inspire future studies on developing diverse pest control methods given the adaptive and neutral genetic differentiation among *D. suzukii* populations. Understanding the extent of local adaptation and its potential environmental drivers will also help predict the spread and future distributions of invasive species (Colautti and Lau 2016). More broadly, the enhanced understanding of how organisms may adapt to geographical, climatic and artificial selective pressures from this study will also be of value in assessing the susceptibility of natural populations to climate change (Kellermann et al. 2012) and human activities (Barange et al. 2010).

## Materials and Methods

### Fly Collection, DNA Preparation, and Pooled Sequencing

Fly samples from 29 populations were used, seven of which were sequenced for the present study. The fly samples sequenced in this study were collected from wild *D. suzukii* populations in two states of the USA, two provinces of Japan, and three European countries (Figure 1; Table S1). Pooled whole adult flies (n = 100 ∼ 183) from each population (Table S1) were used for DNA extraction as previously described (Langley et al. 2011). Library preparations were conducted at the Next Generation Sequencing Core of University of Wisconsin Madison Biotechnology Center (https://dnaseq.biotech.wisc.edu), where pair-end (PE) reads at the length of 150bp were then generated for each of seven pooled DNA samples on an Illumina NovaSeq 6000.

Pool-sequenced reads of 22 additional *D. suzukii* population samples, including from Europe, the Americas, and Asia, were obtained from public data provided by Olazcuaga et al. (2020) at EBI’s SRA (Figure 1; Table S1). Taken together, we formed a comprehensive dataset of 29 populations sampled from native and invasive ranges of *D. suzukii*.

### Quality Control, Alignment, Contamination Analysis, and Variant Calling from Pool-seq Data

To maximize the quality of our analyzed data, we built a high-throughput assembly and quality control pipeline *poolWGS2SNP* with optimized performance, stringent filtering, compatibility with large numbers of genomic contigs, and customized functions to call high-confidence single-nucleotide variants from pool-sequenced data in *D. suzukii* (Figure S6), in part by utilizing resources from the DrosEU bioinformatics pipeline (Kapun et al. 2020).

As an initial quality control of raw PE reads, adapters were removed, and the 3’ end of reads with base quality < 20 were trimmed using *fastp* (Chen et al. 2018). Further trimming was performed using a self-developed python program *filter_PE_length_mem.py* (see Data Availability), where any pair of forward and reverse reads with less than a total of 150 bases with base quality (BQ) ≥ 20, as well as any individual reads with less than 25 bases with BQ ≥ 20 were discarded.

The trimmed and qualified reads were then mapped against the recently released near-chromosome level *D. suzukii* genome assembly Dsuz-WT3_v2.0 that covers autosomes and the X chromosome (Paris et al. 2020) using *bwa mem* (Li 2013). Reads with a mapping quality below 20 were then removed using *Samtools* (Li et al. 2009). We used Picard’s *SortSam* to sort BAM files, and used Picard’s *MarkDuplicates* to mark PCR duplicates to avoid false variant calls (http://broadinstitute.github.io/picard). Indel identification and realignment around indels were performed using GATK’s *RealignerTargetCreator* and *IndelRealigner* (Auwera and O’Connor 2020). Finally, alignments in BAM format were checked for formatting errors using Picard’s *ValidateSamFile*. Summary statistics for quality checking of BAM files were generated using *bamdst* (https://github.com/shiquan/bamdst).

We then checked sample contamination for both newly and previously reported pool-seq data (Table S3), by estimating the proportion of pool-seq reads from different *Drosophila* species as the proportion of reads assigned to species-discriminating *k-mers* (i.e., unique sequences of each species’ reference genome assembly) using the approached described by Gautier (2023). Although the estimation should be reliable, to fully eliminate contaminated reads is not currently practical because substantial proportion of reads can not be confidently assigned to any species (up to 42% of total pool-seq reads), due to the sequence similarity among genomes within the *Drosophila* family (Gautier 2023). Therefore, we focus on identifying samples with contamination and interpreting results these samples with caution.

To call SNPs, we merged the quality-checked BAM files of all population samples into one file using Samtools *mpileup*, only retaining alignments with mapping quality no less than 20 and sites with base quality no less than 20. Variant calling was then performed on the mpileup file using the heuristic SNP caller *PoolSNP* (Kapun et al. 2020). We used a nominally low value for the parameter miss-frac (0.001) to require for each population sample individually, that depth of coverage at a given site be 12 or greater (min-cov = 12), and that this site not be in the top 1% of sites genome-wide for depth of coverage (max-cov = 0.99; calculated separately for each population and for autosomal and X-linked contigs), in order to filter sites subject to copy number variation. In the initial data set used for analysis of genome-wide diversity, we avoided potential biases from allele frequency filters by using min-count = 1 and min-freq = 0. We termed the resulting high-quality sites as ‘analyzed sites’ for brevity.

### Identifying Autosomal and X-linked Contigs

We chose to perform all population genomic analyses and whole-genome scans separately for SNPs from autosomes and the X chromosome for the following reasons: 1) autosomal and X-linked variants have different allelic sample sizes as samples were obtained from both male and female flies; 2) autosomes and the X chromosome could reflect different demographic histories and outcomes of natural selection, *e.g.*, the lower effective population size of the X chromosome than autosomes could lead to a higher impact of bottlenecks and selection on genomic diversity; 3) unbalanced sex ratios and male-biased dispersal could further differentiate autosomal versus X chromosome variation (Clemente et al. 2018; Olazcuaga et al. 2020).

Since the assembly of *D. suzukii* reference genome is still at the contig level, chromosomal identities of each contig are needed to perform separate analyses. However, 497 contigs that represent ∼43% of the assembly length have not been unambiguously mapped onto chromosome arms of the *D. melanogaster* dm6 genome assembly. Although 264 of the unplaced contigs had been assigned to autosomes and the X chromosome based on a female-to-male read depth ratio, 233 contigs that represent ∼5% of the genome remained unassigned due to the lack of statistical power (Paris et al. 2020).

Given our interest in accurately analyzing a larger proportion of the euchromatic genome, we identified ∼70% of these 233 unassigned contigs as autosomal and X-linked based on the correlation between the mean read depth of each contig (among population samples) and that across unambiguously aligned autosomal or X-linked contigs. We chose Spearman’s rank correlation instead of the Pearson correlation, as the distribution of depth data failed the assumption of bivariate normality. A contig that has mean depth significantly correlated with that of either known autosomal or X-linked contigs was assigned to the chromosome with a higher correlation coefficient. Our method completely confirmed all prior mapping-based assignments and had a ∼96% consistency with the previous assignment based on female-to-male read depth ratios. Inconsistent assignments for four contigs were corrected according to our method (Table S4). The eight previously-assigned contigs that could not be assigned using our method, as well as other unassigned contigs using all methods (totaling ∼2.7 mb), were excluded from downstream GEA analyses, because the assignment information is needed for estimating effective sample size that were used to correct allele count data as an input to GEA.

### Annotating Genomic Features and Estimating Divergence

To explore genomic diversity at synonymous sites and selective constraint for other site types in *D. suzukii*, we classified the reference genome into nine exclusive categories of site degeneracy and function (Lange and Pool 2018), including: non-degenerate (i.e., nonsynonymous) sites, two-, three-, and four-fold degenerate (i.e., synonymous) sites, 3’ and 5’ UTRs, RNA-coding genes, introns, and intergenic regions. From input files including the eukaryotic codon table, the published genome sequence and GFF3 annotation obtained at NCBI RefSeq, we generated a letter-coded annotation (in FASTA format) mirroring both strands of the whole genome sequence of *D. suzukii* and a coordinate-based annotation (in BED format) that combines adjacent sites of the same category into a single row. Degeneracy was determined based on the standard codon table. 5’ UTRs were defined as regions between the start of the first exon and the start of the first coding sequence (CDS), while 3’ UTRs were defined as regions between the end of the last exon and the end of the last CDS. In cases of overlapping genes and alternative splicing that raise annotation conflicts, we followed an annotation priority in the category order listed above.

We then estimated the divergence between *D. suzukii* and its close relative *D*. *biarmipes* in each of these categories (Suvorov et al. 2022). We obtained results of multiple sequence alignment between the current reference genomes of *D. suzukii* and *D*. *biarmipes* (Paris et al. 2020). For each site category of *D. suzukii*, the unpolarized divergence was estimated as the number of substitutions over the total number of sites within aligned blocks of reference genome sequences.

### Estimating Nucleotide Diversity, *F_ST_ D_XY_*

To compare genome-wide polymorphism among populations, we estimated nucleotide diversity (*π*) across SNPs at four-fold degenerate sites (*π_S_*) in addition to that at all categories of sites (*π*_=_), as *π*_>_ estimation is relatively less affected by sequencing errors than nucleotide diversity estimated from other site categories (due to a higher ratio of real variation to errors). To calculate *π* for each population sample, we adopted an unbiased estimator of nucleotide diversity (*θ^^^_π_*) based on heterozygosity (*Π*), which has been optimized for pool-seq data (Ferretti et al. 2013). Numerically,

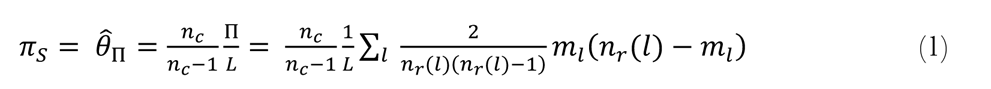

Here, *L* represents the total number of genome-wide analyzed sites. Of a given population sample, *n*_r_(*l*) represents the read depth of the top two alleles at the *l*th site (i.e., SNP) and *m*_L_ represents the minor allele count. *n_c_* as a normalization factor represents the haploid sample size for either autosomes or X chromosome in a pool (Table S1). Strictly speaking, *n_c_* as a normalization factor should represent equally contributing chromosomes in a pool. Nevertheless, for our data it is sufficient to use haploid sample size for either autosomes or X chromosome to approximate *n_c_* in the above equation, as the estimation of *θ^^^_π_*is not substantially affected by the precise value of *n_c_* when the number of individuals in the pool is large. The above formula is a simplified version for SNP data, based on equation 3 in Ferretti et al. (2013).

To examine patterns of polymorphism across chromosome arms, we also estimated window nucleotide diversity (*π*_X_) for all polymorphic sites. Each window was defined as a continuous genomic region that includes 125,000 analyzed sites (Figure 2). Since chromosomal identity was required in this analysis, we only took windows from 32 major contigs that contain at least one full-size window and were unambiguously mappable to a chromosome arm of the *D. melanogaster* dm6 genome assembly. Although such contigs only make up 57% of the *D. suzukii* genome assembly, they contain a relatively larger proportion of all identified SNPs (72%), and thus are still representative of genome-wide polymorphism.

To estimate genome-wide pairwise *F_ST_* between populations, we adopted an unbiased multi-loci estimator known as Reynolds’ estimator of the coancestry coefficient, which accounts for unequal sample sizes among populations and is applicable for more than two alleles at a site (Reynolds et al. 1983). We first heuristically partitioned the genome into windows that exceeded a cross-sample average accumulated heterozygosity threshold of 100. The genome-wide *F_ST_* was then calculated as an average of window-*F_ST_ F_ST_*(*w*) weighted by the number of analyzed sites within each window (*L*(*w*)), where *F_ST_*(*w*) was calculated as a weighted average of single-site ratio estimators. Numerically,

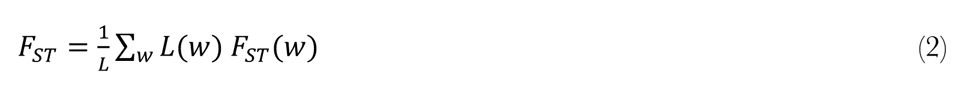

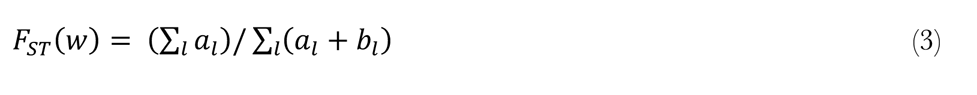

where

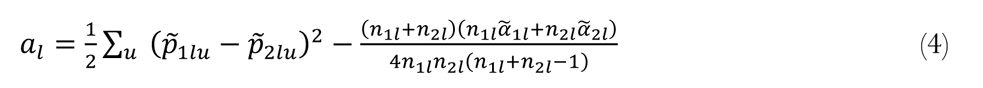

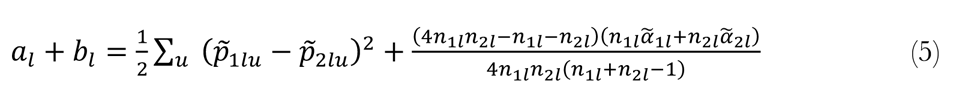

At the *l*th site in each population, *p^~^_1lu_* and *p^~^_2lu_* represent the frequency of the *u*th allele at the *l*th site; *a^~^_1l_* and *a^~^_2l_* represent the heterozygosity; *n_1l_* and *n_2l_* represent the sample size. Unlike the sequencing of individual genomes, pool-seq induces an uncertainty in the number of individual alleles actually sequenced at a locus (i.e., effective sample size), and this uncertainty decreases slowly even at high read depth (Ferretti et al. 2013). Since the sample size is an important parameter for *F_ST_* estimation, we took measures to obtain an estimate of the effective sample size, *n*_jL_, at each given site (Ferretti et al. 2013). Numerically,

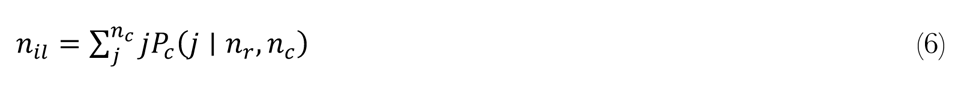

where

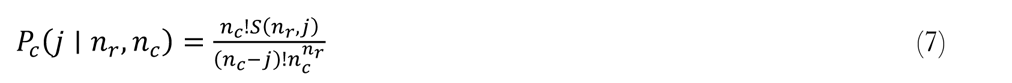

Here, we explicitly estimated the probability of the number of *j* unique lineages sampled at a site given *n*_S_ sampled reads and *n_c_* equally contributing chromosomes in a pool, where *S*(*n*_S_, *j*) are the Stirling numbers of the second kind, defined as the number of ways to partition *n*_S_ reads into *j* non-empty sets (Ferretti et al. 2013). We then estimated *n*_jL_ as the expected number of lineages for each *n*_S_ and *n_c_*. Ideally, *n_c_* should be estimated as 2*n*_*_, where *n*_*_ is the effective pool size representing the number of diploid individuals contributing the same amount of reads to a pool (Gautier et al. 2013; Lange et al. 2022). Although we lack sample replicates to estimate *n*_*_ and therefore used haploid sample size for *n_c_* as an approximation, the probability estimation is still reasonable given that our number of lineages for each pool is large (Table S1) (Ferretti et al. 2013).

Lastly, we estimated genome-wide pairwise *D_XY_* as an absolute measure of population differentiation that is independent of levels of within-population diversity. It was calculated as pairwise differences per site between two populations, divided by *L* total analyzed sites (Nei 1987; Hahn 2018). Numerically,

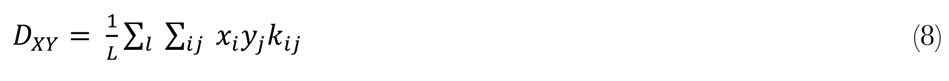

where *x*_j_ and *y*_o_ represent frequencies of the *i*th allele from population *X* and the *j*th allele from population *Y*, and *k*_jo_ is either 1 or 0, depending on whether or not the alleles differ at the *l*th site.

Calculations in this section were all implemented with Python and Shell scripts (see Data Availability).

### Preparing Environmental Data

To generate environmental data for GEA, we selected a preliminary set of 26 candidate environmental variables representing geographic, climatic and land cover-related factors (Figure 3A) that may be relevant in the adaptation process of *D. suzukii* based on prior knowledge (Kellermann et al. 2012; Bogaerts-Márquez et al. 2021). With R packages ‘raster’ (v. 3.5.2) and ‘SpatialPoints’ (v. 1.4.6), we retrieved environmental data of high spatial resolution (∼100 km^M^) in batch for the sampling locations of our 29 populations from online databases WorldClim (Fick and Hijmans 2017) and Esri 2020 Land Cover (Karra et al. 2021). Annual mean values of monthly climatic variables, including mean wind speed, solar radiation, and water vapor pressure, were derived by averaging across 12 months of data.

Due to the large number of statistical tests that would result from running GEA on all the environmental variables one by one, there is an increased difficulty in controlling rates of false discovery. Additionally, including multiple highly correlated variables in a model would lead to multicollinearity issues (Rellstab et al. 2015). To avoid these problems, we calculated a pairwise Pearson correlation matrix from values of environmental factors across sampled locations (Figure 3A; Table S5), and then selected a subset of nine least correlated environmental variables for one-by-one GEA analyses (Figure 3B). To avoid scale inconsistencies between estimated GEA statistics, the environmental differentiation of each population was calculated as the absolute difference between the environmental value of that population and the average across all populations, standardized by the standard deviation (de Villemereuil and Gaggiotti 2015). This standardized differentiation was then input to GEA (Table S5).

### Environmental Association Analyses

To characterize the environmental adaptation of *D. suzukii*, we scanned the whole genome for adaptive loci using the *F_ST_*-based GEA method BayeScEnv (de Villemereuil and Gaggiotti 2015). We chose this specific approach over other GEA methods mainly because it has a lower false positive rate than other GEA approaches in the presence of hierarchical population structure (de Villemereuil et al. 2014; de Villemereuil and Gaggiotti 2015). It also allows for detecting patterns of allele frequency that are not linearly dependent on environmental factors (Rellstab et al. 2015; de Villemereuil and Gaggiotti 2015).

For each environmental variable, the association analyses tested the relationship between environmental and genetic differentiation among populations, for 5,752,156 genome-wide SNPs with a MAF higher than 5%. To control for false positives, we chose stringent model parameters expected to yield extremely conservative results, setting the prior probability of non-neutral models as 0.02 (-pr_jump 0.02) and the prior probability of the competing environment-unrelated locus-specific model as 0.9 (-pr_pref 0.9). These parameters correspond to assumptions that genetic differentiation reflects the action of natural selection in just 2% of the genome, and the focal environmental variable is only expected to be involved at 10% of the non-neutral loci.

To make this GEA analysis computationally feasible with our large SNP set, while still analyzing all qualifying SNPs, we applied a split-run strategy: we subsampled SNPs across concatenated sequences of contigs within the autosomes and the X chromosome separately, and then ran subsamples with BayeScEnv in parallel. Since the null model of population structure is estimated separately in each run, we subsampled non-adjacent SNPs at a fixed interval to limit locus-specific biases in that estimation, where the length of the interval between jointly analyzed SNPs was equal to the total number of subsamples. With a targeted subsample/interval size of up to 10,000 SNPs, we divided the concatenated autosomal contigs into 490 subsamples (with actual subsample sizes of 9982-9983 SNPs), and the concatenated X-linked contigs into 87 subsamples (with actual subsample sizes of 9893-9894 SNPs). Hence, the first autosomal subsample contained SNP #1, SNP #491, and so on.

Convergence of each run was confirmed with the R package ‘CODA’. Individual runs were then merged across autosomes and X chromosome to calculate the genome-wide q-value (*q*) of locally-estimated posterior error probability (*PEP*) across all sites, where we targeted a false discovery rate (*FDR*) of 5% by setting the *q* threshold at 0.05 (Storey 2003; Muller et al. 2006). For downstream analyses, to remove redundancy due to linkage disequilibrium, we pared down closely linked candidate sites by maintaining the site with the lowest *q* within each 20 kb genomic window. To assess the relative levels of support for associations between SNPs and a given environmental variable, we ranked all candidate loci first by *q* and then by the estimated *g* parameter as a tie-breaker, which measures the sensitivity of a locus to environmental differentiation.

### Identifying Candidate Genes

For each candidate SNP, the closest gene in each direction within a 200-exon flanking region that overlapped with it was considered to be associated with that variant, in order to encompass both potential coding and regulatory adaptation. To facilitate clear comparisons among environmental variables with different numbers of significant variants, we focused on the top 500 candidate genes that were linked to variants with the lowest significant *q* and highest g within each environmental variable (Table S7).

### GO Enrichment and Semantic Clustering

Gene ontology (GO) enrichment of the top 500 candidate genes associated with candidate SNPs was performed via genomic permutation of outlier SNP positions (100,000,000 replicates), which accounts for the variability of gene length and the clustering of functionally related genes, as described in previous work (Pool et al. 2017). For each GO category, a p-value indicated the proportion of permutation replicates in which an equal or greater number of genes was implicated.

We then prioritized the most informative and significant GO terms and removed redundant terms that potentially share similar groups of genes by clustering GO terms based on their semantic similarity and ranking representative terms of each cluster by their p-value (Reijnders and Waterhouse 2021). For GO terms that were shared among associations with multiple environmental variables, a combined p-value was calculated from the p-values of independent enrichment tests using Fisher’s method (Fisher 1938).

## Author contributions

SF, JEP, SDS, and CG designed the research. SPD collected data for the project. SF analyzed the data. SF, JEP, and SDS wrote the paper. SDS, JEP, and CG obtained funding for the research.

## Acknowledgments

We thank members of the Pool lab for helpful comments on this manuscript. We also thank Masahito Kimura, Samantha Tochen, and Carandale Farms for assistance with fly collection. We also thank Mathilde Paris for providing a multiple alignment file across *Drosophila* species and updated genomic annotations, as well as Martin Kapun for assistance with SNP calling, and Pierre de Villemereuil for helping with our GEA analyses. The UW-Madison Center for High Throughput Computing provided computational assistance and resources for this work.

## Data Availability

All sequence data generated for this project are available from the NIH Short Read Archive under project PRJNA973110, with specific sample information given in Table S1, Supplementary Material online. All computational scripts created for this study have been uploaded to https://github.com/Sfeng666/poolWGS2SNP (for WGS data processing and variant calling) and https://github.com/Sfeng666/Dsuz_popgen_GEA (for population genetics analyses and GEA).

## Supplementary Material

Supplementary figures S1–S6 and tables S1–S7 are available at Evolution online.

