## Supplemental Figures for "Genomic Diversity Illuminates the Environmental Adaptation of *Drosophila suzukii*"

### Supplementary Figures

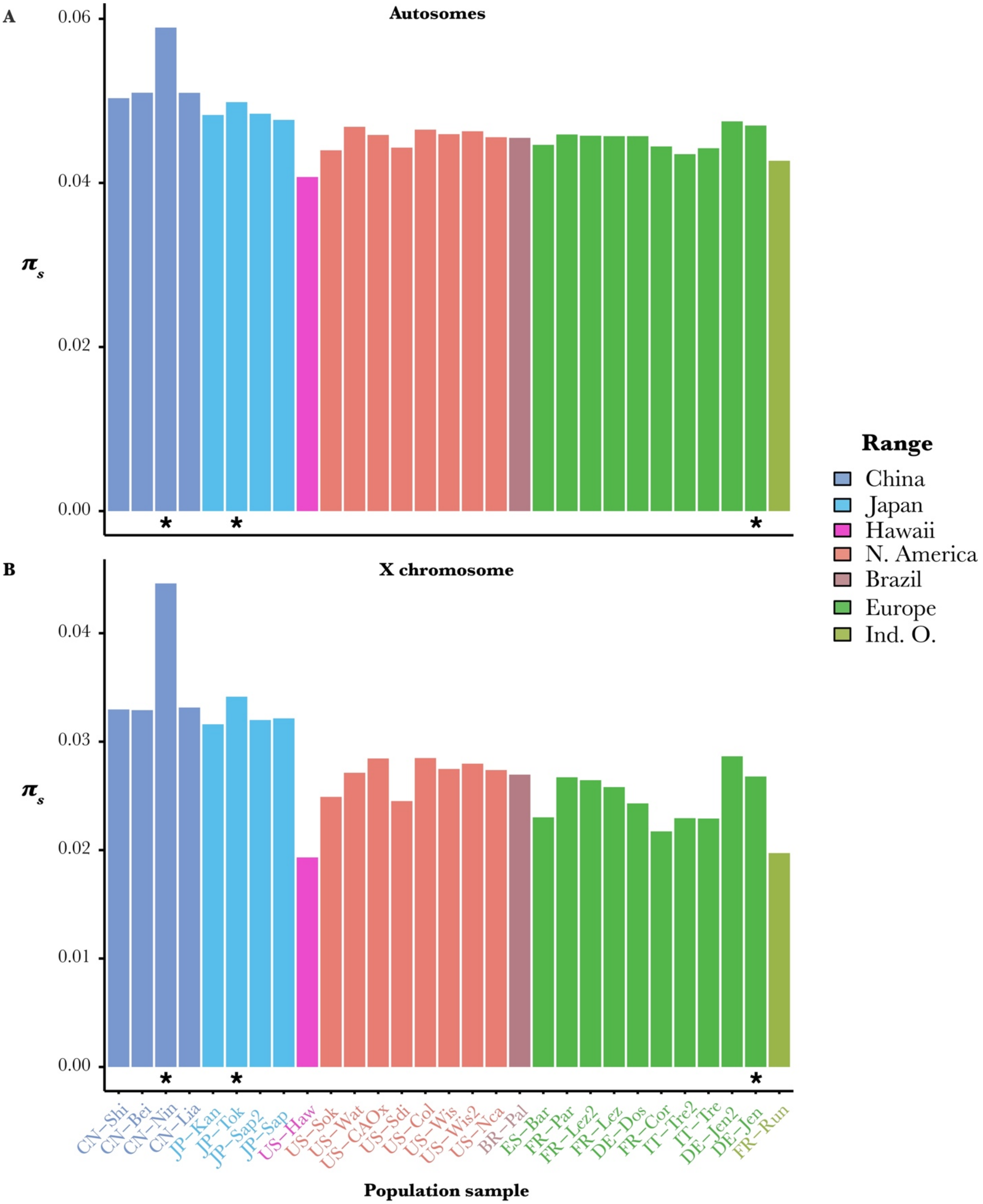


**Figure S1.** Synonymous nucleotide diversity implies invasive bottlenecks of introduced populations. $\pi_s$ was calculated across SNPs in (A) the autosomes and (B) the X chromosome. Populations are colored by their geographical ranges. Asterisks indicate samples contaminated by other *Drosophila* species, which may affect estimation of $\pi$.

A

B

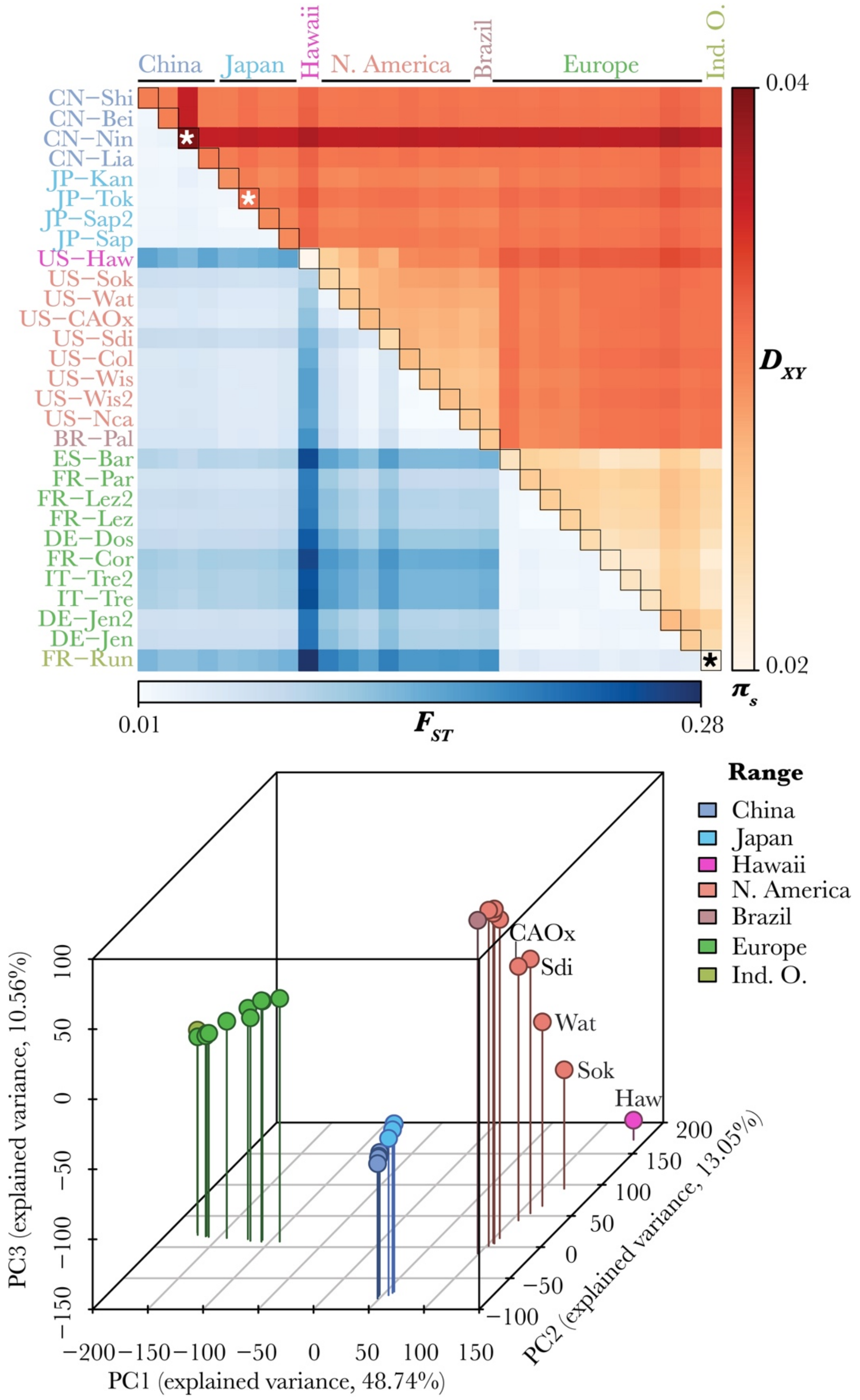


**Figure S2.** X chromosomal polymorphisms recapitulate continent-level genetic structure and maximal diversity in eastern China. (A) X chromosome-wide pairwise $F_{ST}$ (lower triangle), $D_{XY}$ (higher triangle), and $\pi$ (diagonal) across synonymous SNPs are displayed as a heatmap. Population names are colored by their geographical region. Asterisks indicate samples contaminated by other *Drosophila* species, which may affect estimation of $\pi$ and $D_{XY}$. (B) X chromosomal genetic structure is shown by three-dimensional PCA based on allele frequencies of the two most frequent alleles across all populations. Each dot represents a population. Labeled are Hawaii and western coastal US populations.

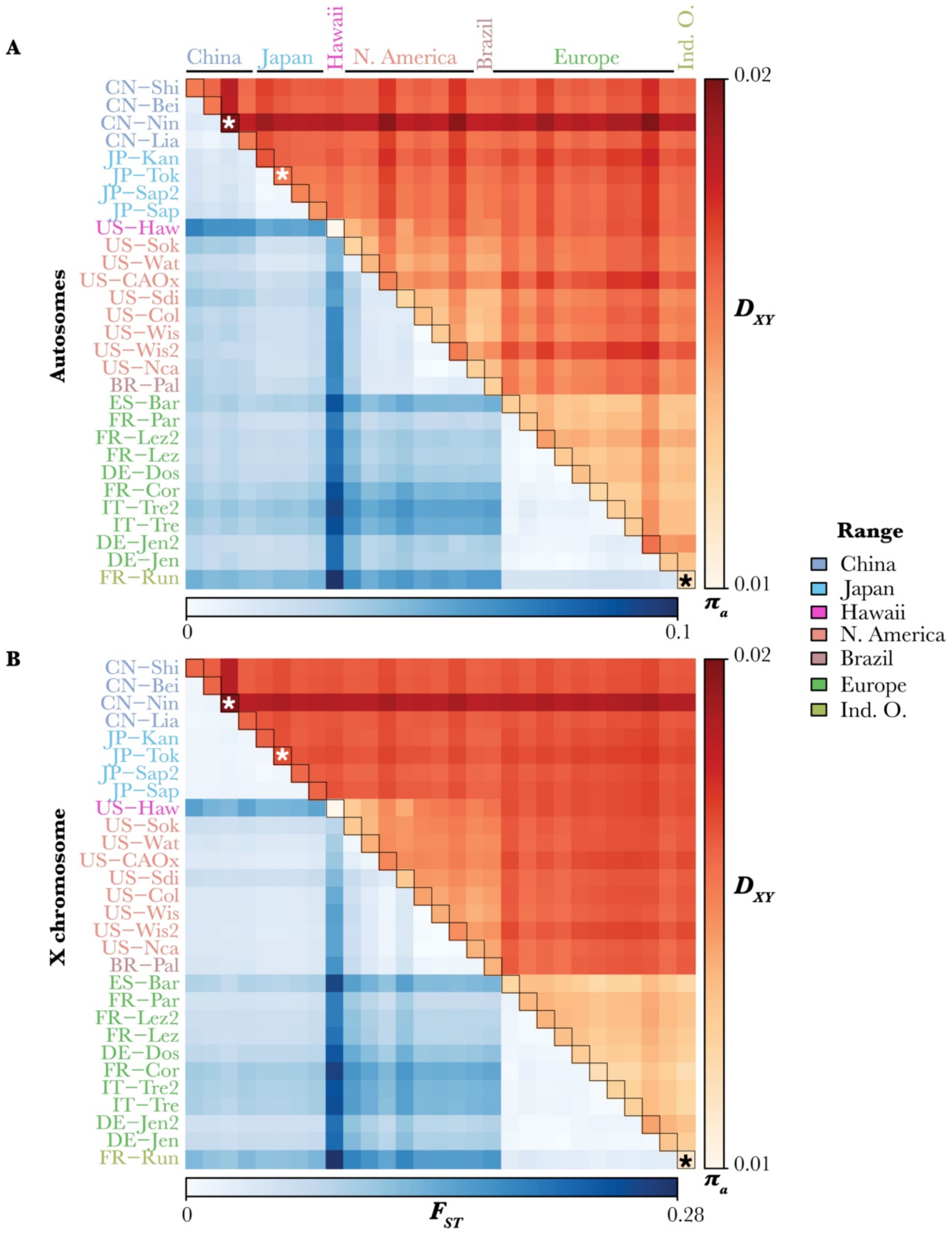


**Figure S3.** Genome-wide polymorphism at all sites recapitulates continent-level genetic structure and maximal diversity in eastern China. (A) Autosome- and (B) X chromosome-wide pairwise $F_{ST}$ (lower triangle), $D_{XY}$ (higher triangle), and $\pi$ (diagonal) across all SNPs are displayed as a heatmap. Population names are colored by their geographical region. Asterisks indicate samples contaminated by other *Drosophila* species, which may affect estimation of $\pi$ and $D_{XY}$.

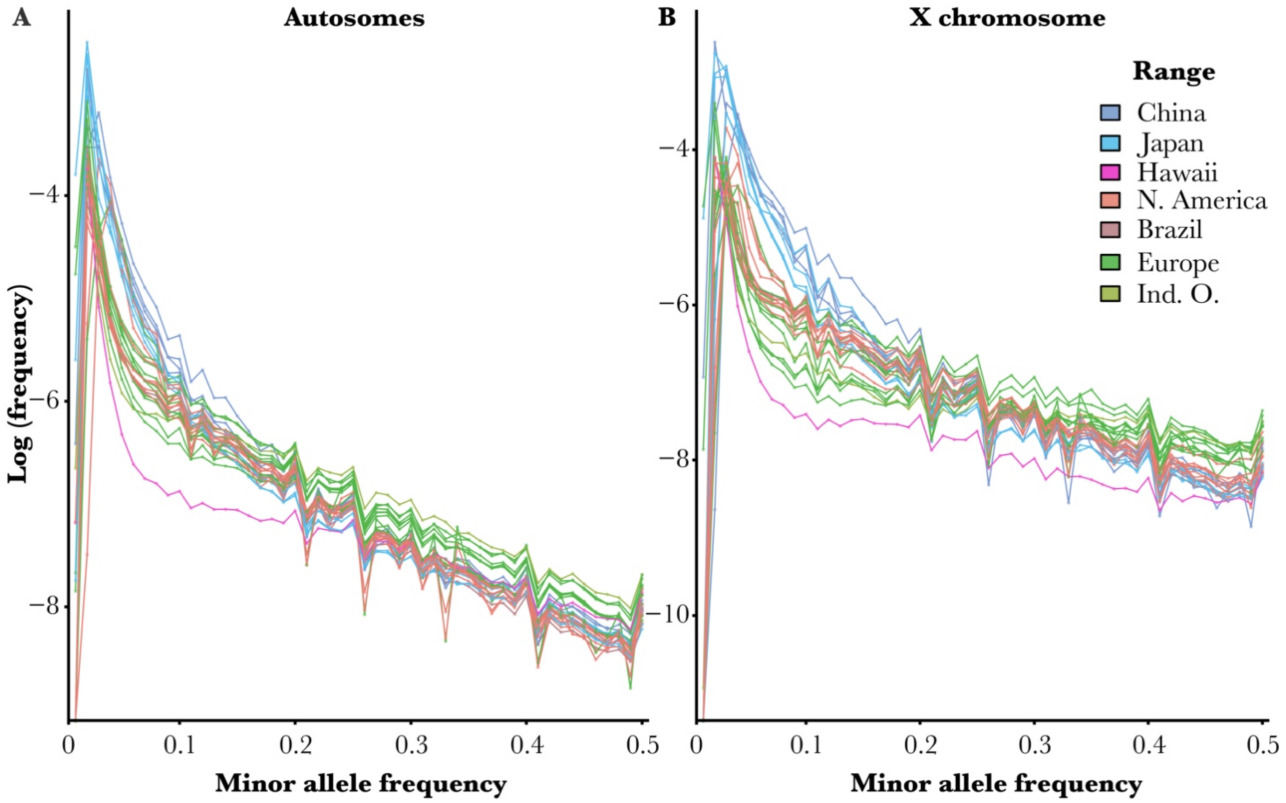


**Figure S4.** Allele frequency spectra of SNPs across (A) autosomes and (B) the X chromosome show a greater loss of rare alleles in invasive populations. Each line represents a population sample, colored by its geographical range. The spectrum is depicted as a distribution of the natural logarithm-transformed frequency of minor alleles within each minor allele frequency (MAF) interval (width = 0.01) from zero to 0.5, where the intervals are closed on the right. Invariant sites are excluded.

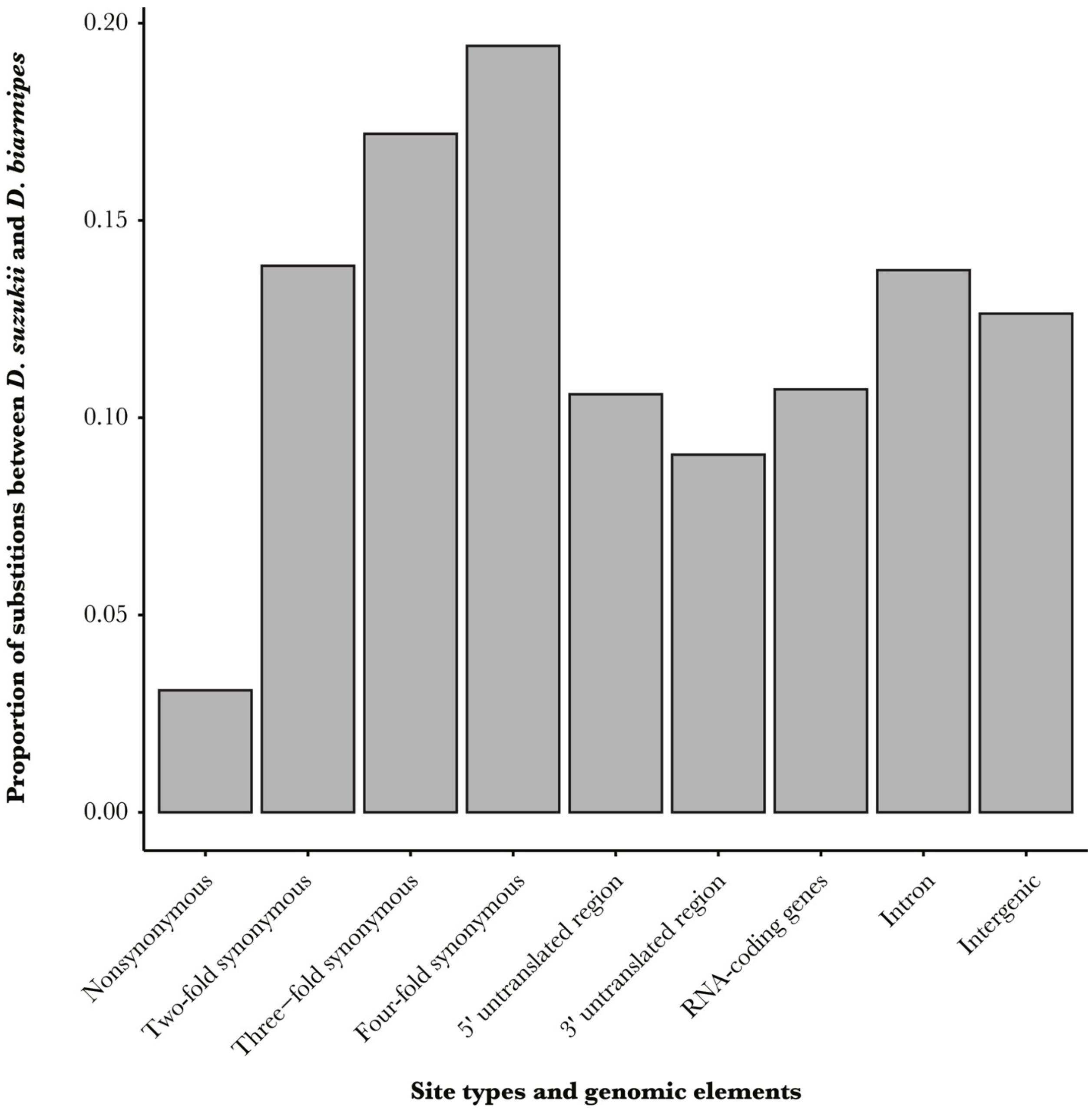


**Figure S5.** The divergence of site types and genomic elements indicates levels of selective constraint. For each site category, the divergence between *D. suzukii* and *D. biarmipes* was estimated as the number of substitutions to the total number of sites within aligned blocks of reference genome sequences.

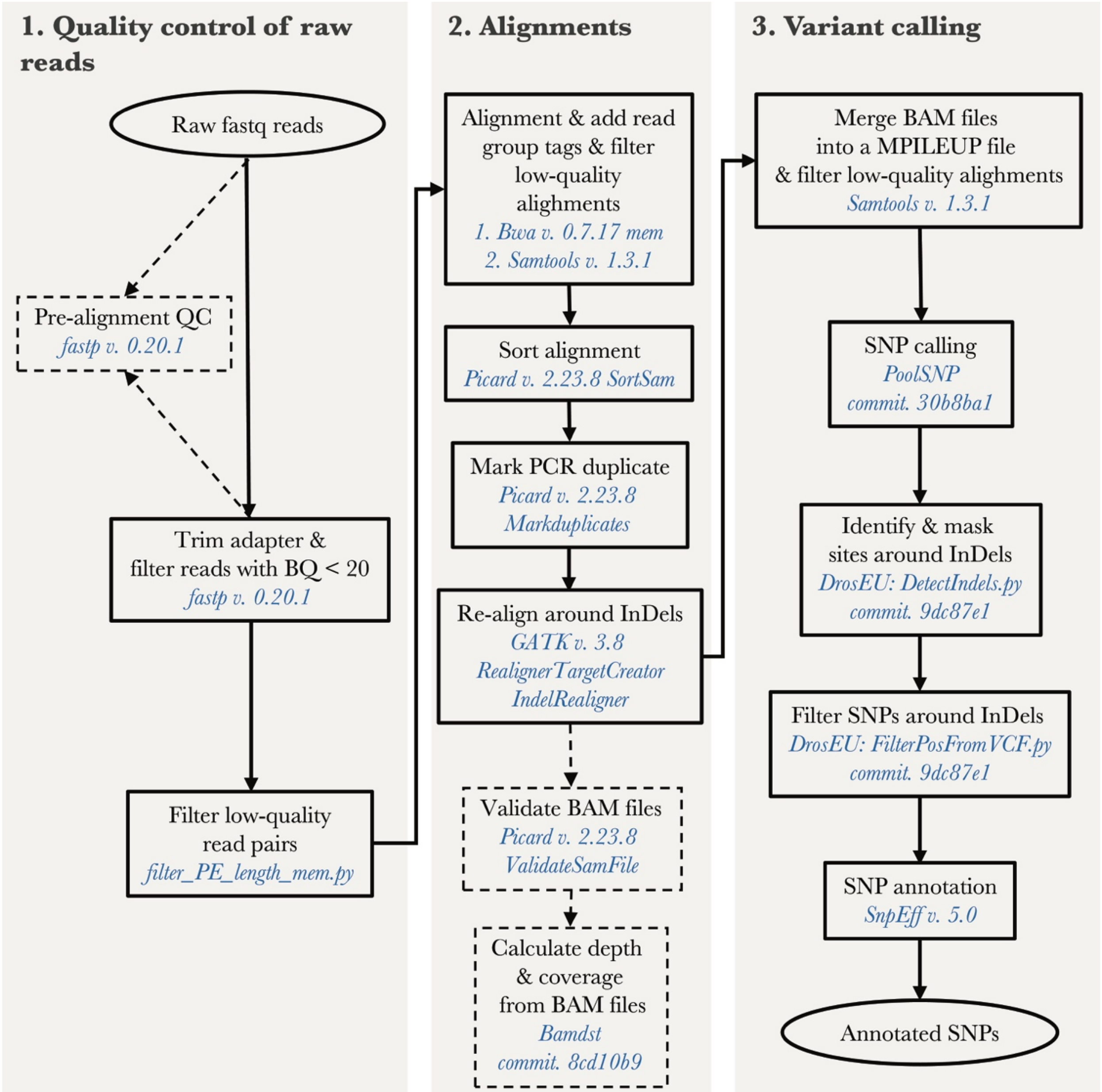


**Figure S6.** The *poolWGS2SNP* analysis pipeline to call SNPs from pool-seq raw reads. Grey shading and bold text represent the three major parts of this pipeline. Input and output are indicated by elliptical boxes. Required steps are indicated by rectangular boxes and arrows in solid lines. Optional steps are indicated by dashed boxes and arrows. Names and versions of used software are colored in blue. Code is available at https://github.com/Sfeng666/poolWGS2SNP.
